# Skip and bin: pervasive alternative splicing triggers degradation by nuclear RNA surveillance in fission yeast

**DOI:** 10.1101/010033

**Authors:** Danny A. Bitton, Sophie R. Atkinson, Charalampos Rallis, Graeme C. Smith, David A. Ellis, Daniel C. Jeffares, Yuan Y.C. Chen, Michal Malecki, Sandra Codlin, Michal Lubas, Jean-François Lemay, François Bachand, Cristina Cotobal, Samuel Marguerat, Juan Mata, Torben Heick Jensen, Jürg Bähler

## Abstract

Exon-skipping is considered a principal mechanism by which eukaryotic cells expand their transcriptome and proteome repertoires, creating different slice varaiants with distinct cellular functions. Here we analyze RNA-seq data from 116 transcriptomes in fission yeast (*Schizosaccharomyces pombe*), covering multiple physiological conditions as well as transcriptional and RNA processing mutants. We applied brute-force algorithms to detect all possible exon-skipping events, which were ubiquitous but rare compared to canonical splicing events. Exon-skipping events increased in cells deficient for the nuclear exosome or the 5’-3’ exonuclease Dhp1, and also at late stages of meiotic differentiation when nuclear-exosome transcripts were down-regulated. The pervasive exon-skipping transcripts were stochastic, did not increase in specific physiological conditions, and were mostly present at below 1 copy per cell, even in the absence of nuclear RNA surveillance and late during meiosis. These exon-skipping transcripts are therefore unlikely to be functional and may reflect splicing errors that are actively removed by nuclear RNA surveillance. The average splicing error-rate was ∼0.24% in wild-type and ∼1.75% in nuclear exonuclease mutants. Using an exhaustive search algorithm, we also uncovered thousands of previously unknown splice sites, indicating pervasive splicing, yet most of these novel splicing events were rare and targeted for nuclear degradation. Analysis of human transcriptomes revealed similar, albeit much weaker trends for pervasive exon-skipping transcripts, some of which being degraded by the nuclear exosome. This study highlights widespread, but low frequency alternative splicing which is targeted by nuclear RNA surveillance.

## Introduction

Splicing is a fundamental step in gene expression which determines the information content of messenger RNAs (mRNAs). It is a highly accurate process that relies on the ability of the spliceosome to identify sequence signals and remove non-coding intronic segments from precursor mRNAs (pre-mRNAs) and thus generate translatable mRNAs. In its simplest form, splicing requires several intronic sequence elements for intron excision, including splice donor and acceptor sites, a branch-site and a polypyrimidine tract (Wang and Burge 2008). Another layer of complexity, alternative splicing, results in the production of multiple transcripts from a single gene, some of which may be condition-or tissue-specific (Wang et al. 2008). These transcripts, referred to as alternative splice variants, carry a non-consecutive combination of exons and are thought to be a major source of transcriptome and proteome diversity in multicellular eukaryotes (Demir and Dickson 2005; Keren et al. 2010). Alternative splicing can also modify post-translational modifications sites (Merkin et al. 2012), or affect translation and protein localisation (Stamm et al. 2005). Splicing is tightly integrated with gene regulation (Braunschweig et al. 2013; Bentley 2014), and it can also control gene expression via nonsense-mediated (Le Hir et al. 2003; Guan et al. 2006) or spliceosome-mediated (Volanakis et al. 2013) decay pathways (NMD and SMD, respectively).

Alternative splice variants may arise via single or multiple exon-skipping or intron-retention events or via alternative 5’- or 3’-splice sites (Wang et al. 2008). The extent of alternative splicing in the human transcriptome is potentially immense, with >90% of genes showing more than one isoform (Wang et al. 2008). Alternative splicing can be governed by sequence elements such as intronic and exonic splicing enhancers or silencers, which serve as binding sites for RNA-binding proteins that facilitate or inhibit splicing (Fairbrother et al. 2002; Yeo et al. 2007). A ‘splicing code’, which dictates the final composition of a given transcript in a given condition, remains elusive however. It is now apparent that splice site selection is not only dictated by sequence information but also by intricate interplays between chromatin, RNA polymerase II and the spliceosome (Alexander and Beggs 2010; Schwartz and Ast 2010).

Unspliced and partially spliced transcripts can be deleterious for the cell (Egecioglu et al. 2012; Sayani and Chanfreau 2012), and alternative or aberrant splicing has been implicated in human diseases (Cooper et al. 2009). To deal with this problem, cells use several nuclear and cytoplasmic quality-control pathways, which degrade faulty transcripts. In human and yeast, the nuclear exosome acts as the first and main line of protection (Garneau et al. 2007; Schmid and Jensen 2008; Houseley and Tollervey 2009). The exosome is a multi-protein complex containing nine non-catalytic core subunits, which interact in the nucleus with the two catalytic subunits Rrp6 and Dis3 that confer 3’-5’ exoribonuclease and endoribonuclease activities, respectively (Houseley et al. 2006; Garneau et al. 2007; Schmid and Jensen 2008; Schneider and Tollervey 2013). RNA degradation by the exosome is facilitated by polyadenylation activity of the TRAMP complex (LaCava et al. 2005; Schmidt and Butler 2013), by helicase activity (Houseley et al. 2006; Schmidt and Butler 2013), and by decapping and deadenylation (Garneau et al. 2007; Arribas-Layton et al. 2013). In budding yeast, unspliced and partially spliced transcripts, as well as transcripts with abnormal exon-skipping, are actively degraded by the nuclear exosome, the 5’-3’ nuclear exonuclease Rat1 (Dhp1 in fission yeast) or the RNase-III endonuclease Rnt1 (Egecioglu et al. 2012; Sayani and Chanfreau 2012).

Defective transcripts that evade degradation by nuclear surveillance may still be degraded by cytoplasmic surveillance pathways (Sayani and Chanfreau 2012). In yeast, cytoplasmic degradation also occurs in either 3’-5’ or 5’-3’ direction, via the core exosome associated with Dis3 (Houseley et al. 2006; Schmid and Jensen 2008), via the 5’-3’ cytoplasmic exonuclease Xrn1 (Exo2 in fission yeast) (Wood et al. 2002; Sayani and Chanfreau 2012), or via the Dis3L2 3’-5’ RNA decay pathway (Malecki et al. 2013). Cytoplasmic mRNA degradation by the exosome is initiated by deadenylation of the transcript and assisted by the helicase activity of the SKI complex (Anderson and Parker 1998; Schmid and Jensen 2008). Cytoplasmic degradation can also be triggered by the NMD pathway that recognizes premature stop codons (Sayani et al. 2008; Kawashima et al. 2014) or by the non-stop decay (NSD) pathway that identifies transcripts lacking stop codons (Frischmeyer et al. 2002). Partially spliced transcripts are degraded by the Dbr1-dependent pathway, which recognizes intronic lariat intermediates (Hilleren and Parker 2003).

Alternative splicing is widely thought to greatly augment protein diversity. A recent analysis of human transcriptomes, however, has revealed that most protein-coding genes express only one dominant transcript that contributes to the proteome in multiple tissues (Gonzalez-Porta et al. 2013). In contrast to tissue-specific gene expression programs, alternative splicing is much less conserved and often lineage-specific (Barbosa-Morais et al. 2012; Merkin et al. 2012). Furthermore, stochastic splicing errors can increase transcript isoform diversity (Melamud and Moult 2009; Pickrell et al. 2010). These findings raise the possibility that a considerable fraction of alternative splice variants simply reflect splicing noise without any cellular function, or with functions other than alternative protein production.

To explore this hypothesis, we analyzed transcriptomes under diverse environmental, physiological and genetic perturbations in the fission yeast, *Schizosaccharomyces pombe*. Approximately 47% of the *S. pombe* genes contain annotated introns, and the splicing machinery is conserved (Käufer and Potashkin 2000). Limited evidence suggests alternative splicing via intron retention (Moldon et al. 2008; Duncan and Mata 2014) or exon-skipping (Awan et al. 2013) for a few

*S. pombe* genes, but some exon-skipping events have been attributed to splicing errors (Bitton et al. 2014). We show here that widespread but rare exon-skipping events, and many other unannotated splicing events, mostly result in the production of rare transcripts that are actively degraded via the nuclear exosome and Dhp1. Analysis of human exosome-depleted cells uncovered weaker, but similar trends of unannotated, pervasive exon-skipping that slightly accumulated in cells with compromised RNA degradation. These observations indicate that alternative splicing via exon-skipping in fission yeast, and, to much lesser extent in humans, represents aberrantly spliced transcripts that are largely destined for degradation in the nucleus.

## Results

We analyzed transcriptomes of wild-type *S. pombe* strains under the following physiological and environmental conditions: vegetative growth in minimal and rich media, stationary phase, quiescence upon glucose and nitrogen limitation, heat stress, and meiotic differentiation. We also analyzed transcriptomes of several mutants with defects in the following RNA processing and transcriptional processes/complexes: core and catalytic subunits of the RNA exosome, nuclear and cytoplasmic 5’-3’ exonuclease, TRAMP and SKI complexes, NMD and NSD, mRNA decapping, poly(A)-binding, cytoplasmic deadenylation, RNA splicing and debranching, RNAi, nucleosome remodeling and RNA polymerase II. Including biological repeats, our analysis encompassed RNA-seq data from 116 transcriptomes from different physiological and genetic perturbations, encompassing both published and original data. A detailed overview of this data set is provided in Supplemental Table S1.

### Exon-skipping events in fission yeast are ubiquitous but rare

To identify sequence reads that represent exon-skipping events among the multiple samples analyzed, we generated a database containing all theoretical exon-exon junctions for all protein-coding genes in *S. pombe*. We aligned all sequence reads both against this junction database and against the reference genome, retaining only reads mapping to a single location, with up to 3 mismatches (3,676,121,463 mappable reads in total) (Fig. 1A). For the exon-skipping analysis, we used all exon-exon junction reads for canonical and exon-skipping events and all exon-intron boundary reads (Fig. 1A).

**Figure 1.**
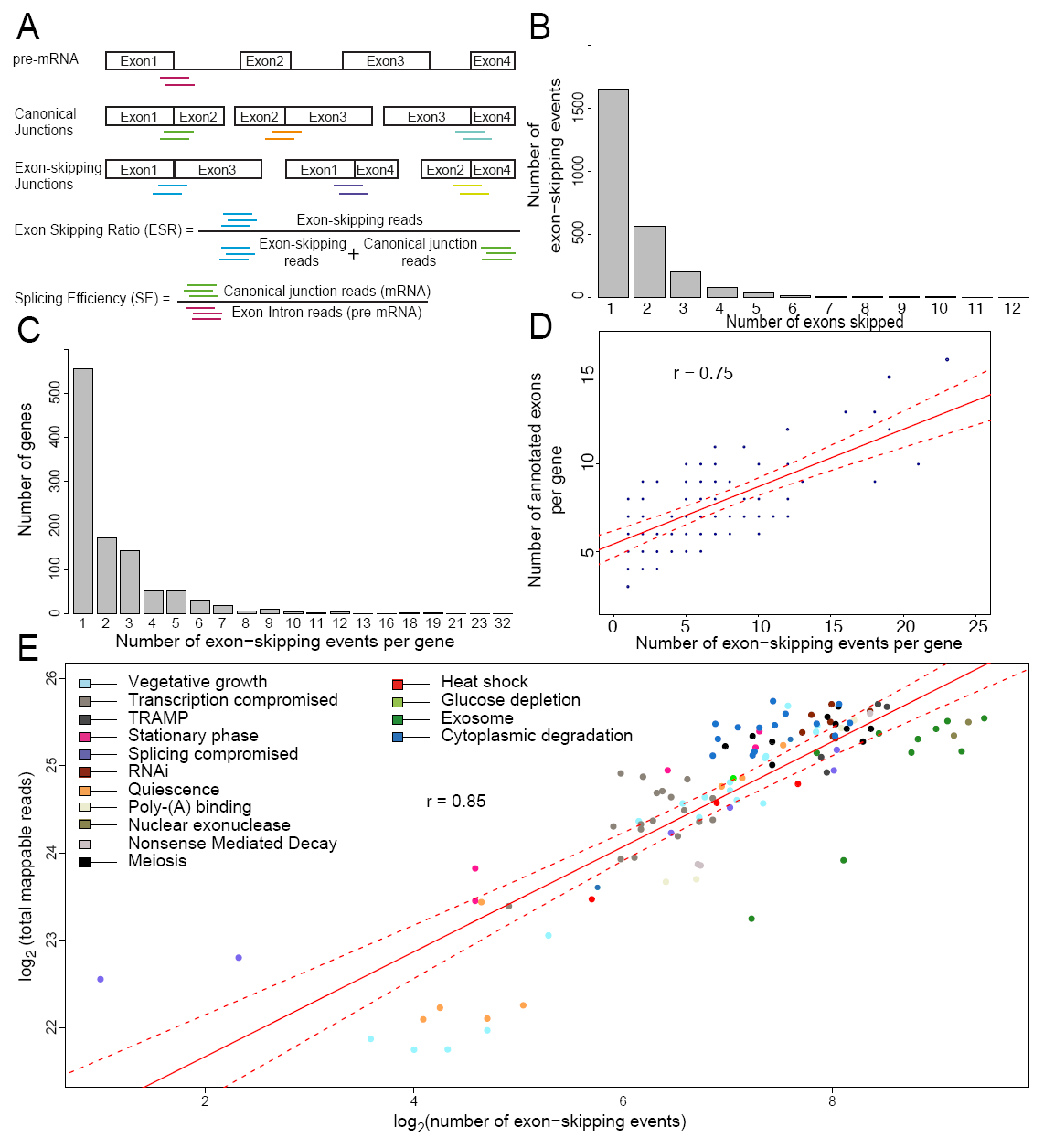
Identification and characterization of exon-skipping events using RNA-seq. (A) Scheme showing pre-mRNA (top) along with all possible canonical and exon-skipping splice isoforms. The colored lines below the junctions represent diagnostic exon-intron reads (pink), canonical exon-exon junction reads (5’-3’; green, orange, cyan), and exon-skipping junction reads (5’-3’; blue, purple, yellow). These diagnostic reads were used to calculate the exon-skipping ratio (ESR), to identify splice isoforms, or to compute the splicing efficiency of a given intron. (B) Total number of exon-skipping events (Y axis) as a function of the number of exons being skipped (X axis; ranging from 1-12 skipped exons during a single event); a total of 2,574 skipping events are shown. (C) Maximum number of exon-skipping events recorded per gene (X axis) as a function of gene number (Y axis). (D) Correlation between numbers of exon-skipping events per gene and numbers of annotated exons in same gene. A total of 1,063 genes were binned based on the number of detected exon-skipping events and the number of their annotated exons; the size of data-points was scaled according to the number of genes in each bin, i.e. normalized by the number of genes in the genome containing the corresponding number of exons; r: Pearson’s correlation coefficient; red line – fitted regression; dotted red lines: 0.95 confidence levels. (E) Correlation between sequencing depth and number of exon-skipping events. Strains were grouped and color-coded according to their cellular function or the condition tested. For full description of each strain and group, see Supplemental Table S1; r: Pearson’s correlation coefficient; red line: fitted regression; dotted red lines: 0.95 confidence levels.

About 2.24% of the mappable reads originated from exon-exon junctions (82,232,577 reads; Supplemental Fig. S1), and only 0.001% of the mappable reads were diagnostic for exon-skipping events (44,616 reads). We initially considered all ‘skipping-reads’ that represent single or multiple exon-skipping events in a given transcript and condition. This analysis identified 2,574 distinct exon-skipping events in 1,063 genes (Supplemental Table S2). Since the fission yeast genome contains only 1,375 genes with at least annotated 3 exons, our finding implies that ∼77.3% of these genes are alternatively spliced by exon-skipping.

The majority of these exon-skipping events only involved the skipping of one exon, and the frequency of exon-skipping decreased with increasing numbers of skipped exons (Fig. 1B). Moreover, genes typically only contained one exon-skipping event, and the frequency greatly decreased with increasing numbers of exon-skipping events per gene (Fig. 1C). The number of exon-skipping events per gene was correlated with the number of exons (Fig. 1D); hence, the likelihood of exon-skipping increased with higher numbers of introns to be spliced. However, the number of exon-skipping events did only marginally correlate with the length of introns or the length of the skipped exons (Supplemental Fig. S2), and with the expression levels of the corresponding transcripts (Supplemental Fig. S3-S4). On the other hand, the number of exon-skipping events strongly increased with sequencing depth (Fig. 1E). This finding reflects that transcripts carrying exon-skipping information are rare, and their identification therefore heavily depends on sequencing depth. Taken together, we conclude that exon-skipping events are infrequent, yet pervasive in the fission yeast transcriptome. The frequency of exon-skipping per gene increases with increasing exon numbers, but is independent of expression level and intron or exon length. These findings are consistent with exon-skipping events largely representing splicing errors.

### Exon-skipping increases in nuclear RNA degradation mutants and during meiotic differentiation

To further investigate the possibility that exon-skipping events represent splicing errors, we calculated the global exon-skipping ratio (ESR), which reflects the proportion of all skipping-reads among all exon-exon junction reads for each sample (Fig. 1A). This analysis revealed a significant enrichment of exon-skipping in the nuclear exosome mutant *rrp6* (Fig. 2A; p <2.2e-16, Cochran-Mantel-Haenszel test, Bonferroni corrected), and in the nuclear exonuclease mutant *dhp1* (Fig. 2A; p <2.2e-16). Also other exosome subunit mutants (*rrp41 and dis3*), components of both nuclear and cytoplasmic exosome complexes, exhibited a pronounced increase in exon-skipping frequency. As expected, strains with a mutated *dis3* RNaseII domain (*dis3_54-ts*) showed a significant but more moderate increase in exon-skipping compared to *dis3, rrp6* or *rrp41* mutants, in line with the documented reduction in exonuclease activity in *dis3_54-ts* cells (Murakami et al. 2007). Note that in the case of *dis3* and *rrp41* mutants, it is practically impossible to determine whether a faulty nuclear or cytoplasmic exosome gave rise to the accumulation of exon-skipping events. However, given the dramatic increase in exon-skipping frequency when the function of known nuclear proteins was impaired (Dhp1 and Rrp6) and the lack of accumulation of exon-skipping events in numerous cytoplasmic degradation mutants, it seems most likely that impaired nuclear exosome activity led to the observed increase in *dis3* and *rrp41* mutants.

**Figure 2.**
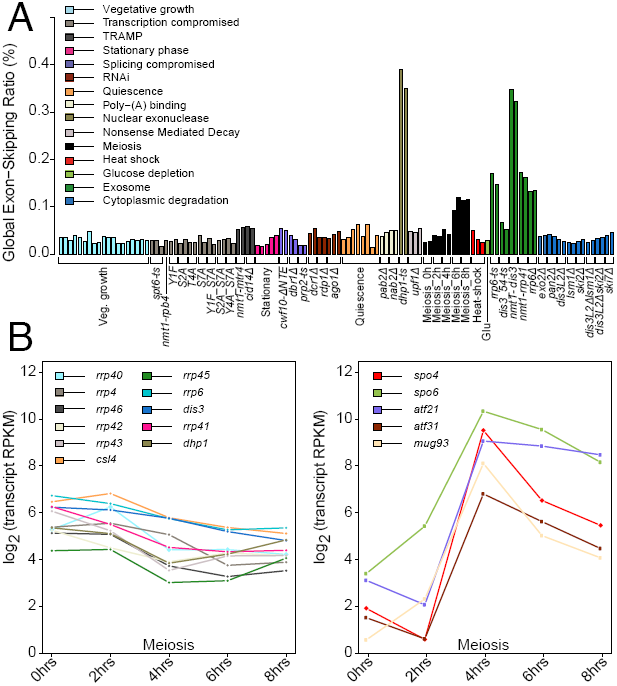
Exon-skipping is rare but accumulates in nuclear surveillance mutants and meiosis. (A) Sample-specific global exon-skipping ratio (ESR) that reflects the proportion of exon-skipping reads among total exon-exon junction reads. Strains as indicated below the bars were grouped and color-coded according to their cellular function or the condition tested. For full description of each strain, see Supplemental Table S1. (B) RNA-seq expression profiles of selected transcripts. Left panel: exosome subunit and nuclear exonuclease transcripts as a function of meiotic progression. Right panel: selected transcripts known to fluctuate during meiosis (RPKM-Reads Per Kilobase per Million; log_2_ scale). Mean expression of two biological replicates is shown at each time-point. The decrease in nuclear-exosome transcripts is significant (p-adjust <0.05 determined by DESeq; Anders and Huber 2010).

More modest, but significant, increases in exon-skipping were also evident in some of the other mutants and conditions tested (Fig. 2A), e.g. mutants with impaired TRAMP complex (*nmt1-mtr4, cid14*Δ*; p <*1.3e-13), or with defective splicing (*cwf10-*Δ*NTE; p <*2.0e-12). Notably, mutants with impaired cytoplasmic degradation showed no significant increase in exon-skipping (Fig. 2A), and the NMD pathway mutant showed only a subtle increase (*upf1*Δ*; p <*1.1e-11). These results indicate that the majority of exon-skipping transcripts are degraded by the nuclear exosome and the Dhp1 exonuclease, while other pathways might play minor or backup roles.

Exon-skipping events also significantly increased during late stages of meiotic differentiation, 6-8 hrs after induction of *pat1-114-*driven meiosis (Fig. 2A; p <2.2e-16), corresponding to late meiosis-II and sporulation (Mata et al. 2002). Notably, many of the core and catalytic nuclear-exosome transcripts significantly decreased during meiosis, particularly at 4 hrs following meiotic induction, with a mean decrease of ∼2.7-fold (Fig. 2B). Approximately a 2-fold decrease was maintained in several exosome subunits also at 6 and 8 hrs following induction. In contrast, only a small fraction of splicing factors showed ≥2 fold decrease in expression during the meiotic timecourse relative to timepoint 0 (∼6-10.5% of 142 genes with RNA splicing and RNA splicing regulation GO categories, GO0008380 and GO0043484, respectively). This finding suggests that the increased exon-skipping frequency in late meiosis could result from lower nuclear surveillance activity.

Taken together, our analysis reveals that exon-skipping events increase most when nuclear RNA degradation pathways are compromised and during late stages of sexual differentiation when transcript levels for components of the surveillance machinery decrease. These results further suggest that most exon-skipping events represent aberrantly spliced transcripts that are actively degraded by nuclear RNA surveillance.

### Exon-skipping variants are stochastic and do not increase in specific physiological conditions

The global ESR revealed overall enrichment of exon-skipping events in specific samples (Fig. 2A). However, this global analysis would not uncover any alternatively spliced variants that may increase in relative abundance in specific samples, i.e. potential isoform switching. To find such transcripts, we computed the local, variant-specific ESR (Fig. 1A; Supplemental Table S3). The local ESR represents the proportion of exon-skipping events among all splicing events at a given locus. As for the global ESR, this analysis showed that exon-skipping was more pronounced in nuclear degradation mutants and during late meiotic stages (Fig. 3A). However, the local ESR was typically close to 0 (average local ESR across entire dataset: ∼0.0037), and was often variable between biological replicates (Fig. 3A; Supplemental Table S3). These results are consistent with exon-skipping events reflecting random splicing errors.

**Figure 3.**
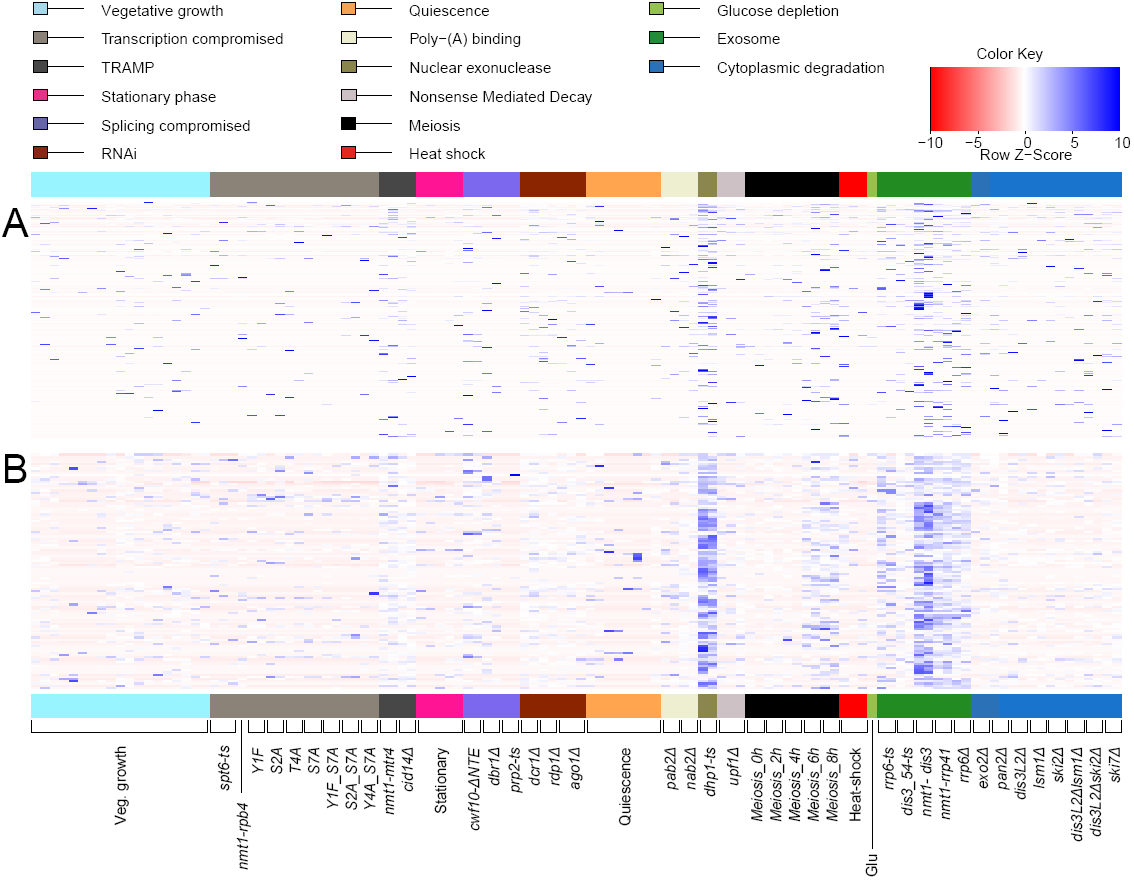
Splice variant-specific, local exon-skipping ratios (ESR) highlighting stochastic nature of exon-skipping. (A) Heatmap representing 2,504 exon-skipping events for which both skipping and canonical reads were identified in at least one sample (actual ratios provided in Supplemental Table S3). Strains as indicated below the colored bar were grouped and color-coded according to cellular functions or conditions. Maximum distance between rows (exon-skipping events) was determined using the “dist” function in R (method “maximum”), followed by hierarchical clustering using the “hclust” function. Row z-score: ratios in each row were scaled by subtracting the mean of the row from each value, followed by division of the resultant values by the SD of the row, i.e. (local ESR value - row mean)/row SD. (B) As in (A) but showing local ESR for high confidence set (111 exon-skipping events supported by at least 9 exon-skipping reads in at least one sample). Only 107 exon-skipping events for which both exon-skipping and canonical reads were identified in at least one sample are represented here (Supplemental Table S4).

RNA-seq experiments are known to suffer from Poisson noise (or counting/shot noise) when read counts are very low (Anders and Huber 2010). Since we initially considered any skipping-read as evidence for an exon-skipping event, it is possible that the observed variability reflects Poisson noise. Therefore, to mitigate the effect of low-read counts, we applied a series of sample-specific skipping-read cutoffs when considering a putative exon-skipping event. As expected, the number of putative exon-skipping events decreased with increasing skipping-read counts (Supplemental Fig. S5A). Notably, ∼51% of the skipping-reads in our dataset (22,817/44,616) originated from only 111 exon-skipping events (∼4.3% of total exon-skipping events), occurring in only 108 genes (Supplemental Table S4). These events were supported by >8 skipping-reads in at least one sample. This observation suggests that a minority of genes are much more prone to exon-skipping than others, raising the possibility that these genes are truly alternatively spliced. As the analyses below show, however, even these higher frequencies of exon-skipping for a sub-set of transcripts most likely represent splicing errors.

The local ESR appeared less variable when only considering the high-confidence set of 111 exon-skipping events (Fig. 3B). But even among this subset, however, the local ESR only showed reproducibly higher values in cells with compromised RNA degradation and during late meiotic stages (Fig. 3B). Moreover, no significant functional enrichment was evident, neither for the entire set of 1,063 genes in which exon-skipping occurred nor for the high-confidence subset of 108 genes. We further ranked the high-confidence set of exon-skipping events based on their local ESR and their reproducibility across the different physiological and environmental conditions tested, excluding genetically perturbed samples (ESR ≥0.1, followed by ranking based on numbers of samples that met threshold in 46/116 samples: vegetative growth, stationary phase, quiescence, glucose depletion, heat-shock, and meiosis). Only 8 exon-skipping events displayed an ESR ≥0.1 in at least 9 of the 46 samples (Supplemental Table S4). The local ESR of these highest-abundance events was still further increased in nuclear RNA surveillance mutants, however, suggesting that even these most frequent exon-skipping events are subject to degradation.

Across the entire dataset, only 904 of 2,574 skipped exons were divisible by 3 with respect to nucleotide number (i.e. did not change the reading frame); this subset did not exhibit any higher number of skipping-reads (∼35%; p >0.05 Supplemental Fig. S5B). Also, the high-confidence set of 111 exon-skipping events was not enriched for skipped exons that were divisible by 3 (34%), and the 8 most frequent exon-skipping events also displayed a similar pattern close to random expectation (3/8; 37.5%). Thus, most exon-skipping transcripts are unlikely to produce a functional protein if the same start codon is being used (Magen and Ast 2005). Taken together, we conclude that exon-skipping in fission yeast is largely random, infrequent and unlikely to be functional in any of the physiological conditions analyzed.

### RNA surveillance increases splicing accuracy

Assuming that all exon-skipping events represent aberrantly spliced transcripts under the conditions tested here, the local ESR reflects the splicing error-rate at a given locus. The global ESR, on the other hand, underestimates the global splicing error-rate, because it was computed using all exon-exon junction reads, including those originating from genes with a single intron. Using the 2,504 exon-skipping events in the 1,049 genes where both canonical and exon-skipping-reads were identified in at least one sample, we computed the mean of the local ESRs for all genes (Supplemental Table S3). These values thus estimate the average splicing error rate at a given locus under each condition (Fig. 4A; Supplemental Table S5).

**Figure 4.**
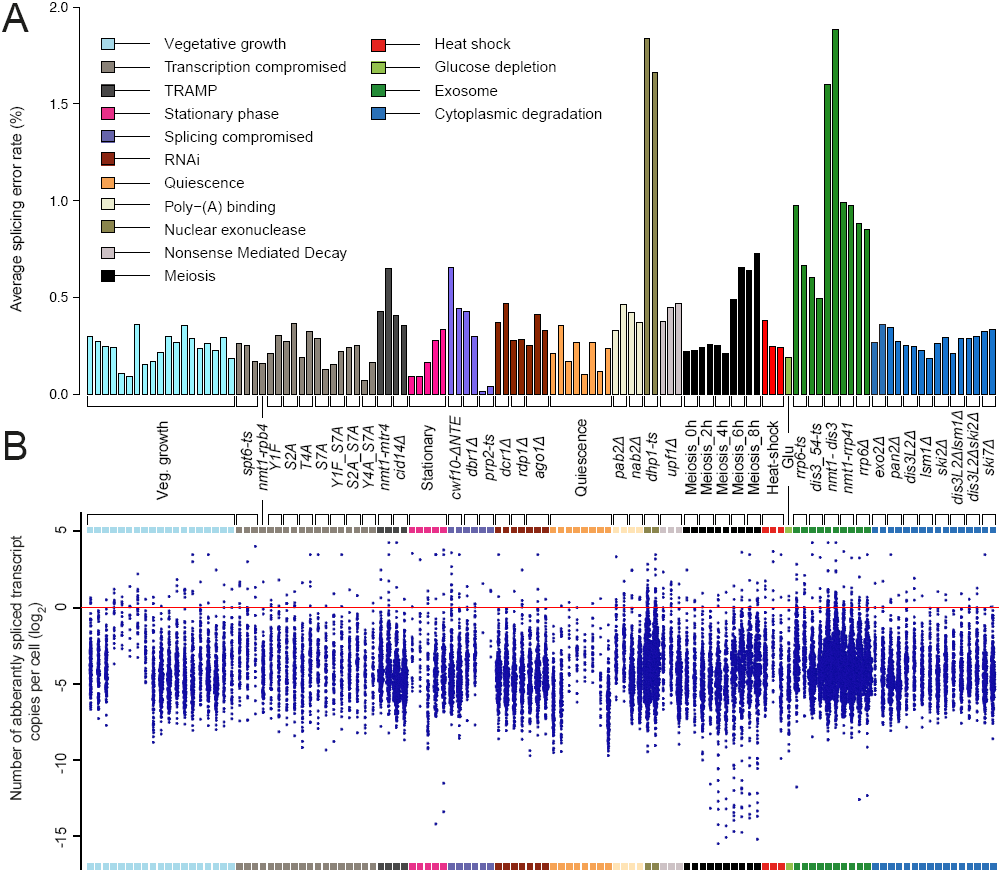
Splice-site selection errors are masked by nuclear RNA surveillance. (A) Average splicing error for a given sample. Set of 2,504 exon-skipping events for which both canonical and non-canonical skipping-reads were identified in at least one sample. The splicing error-rate for a locus is equal to its corresponding local ESR (Supplemental Table S3). The actual values shown here are the averages of all local ESRs in a sample (for actual values, see Supplemental Table S5). Strains as indicated below bars were grouped and color-coded according to cellular functions or conditions. For full description of each strain, see Supplemental Table S1. (B) Number of aberrantly spliced transcripts per cell. To estimate this number, we multiplied the local ESR for each exon-skipping event by its corresponding transcript copy number (Marguerat et al. 2012); for actual values, see Supplemental Tables S6. Absolute numbers of transcripts per cell from proliferating cells were used as gold standard for all samples, except for quiescence where the reported absolute numbers were used. Each dark blue point represents the copy number of aberrantly spliced transcripts for a given gene (log_2_ scale). To avoid log_2_ of 0, exon-skipping events with local ESR/splicing error of 0 were removed. Strains as indicated in (A).

In wild-type cells with intact nuclear RNA surveillance, only ∼0.24% of splicing errors were detectable, lower than estimated for human cells (∼0.7%; Pickrell et al. 2010). With compromised nuclear exonuclease activity (*dhp1-ts*), however, the mean splicing-error rate increased by more than 7-fold (∼1.75%). Strains with compromised Dis3 function (*nmt1-dis3*) displayed a similar average splicing error (∼1.74%), while strains compromised for Rrp6 (*rrp6-ts* and *rrp6*Δ) showed a more moderate increase (∼0.84%). Naturally, these are minimal estimates as the error rate might further increase in the simultaneous absence of several RNA surveillance pathways. An increased splicing error-rate was also observed during late meiotic differentiation (∼0.63%). We conclude that most splicing errors are masked by nuclear RNA surveillance.

### Exon-skipping transcripts are present at below 1 copy per cell

Absolute numbers of transcript copies per cell for all *S. pombe* genes during cell proliferation and quiescence are available (Marguerat et al. 2012). Here we also analyzed the same transcriptome samples (vegetative growth and quiescence; Supplemental Table S1). Assuming that exon-skipping events represent errors in splice-site selection, we used these quantitative transcript data for growing and quiescent cells (Marguerat et al. 2012), along with the local ESRs, to estimate cellular copy numbers for exon-skipping transcripts for each condition and splice varian (Supplemental Table S6).

Exon-skipping transcripts were largely present at below 1 copy per cell, suggesting that these events occur only in a portion of cells in a population (Fig. 4B). Notably, although the diversity of exon-skipping events increased in nuclear RNA surveillance mutants and during late meiotic differentiation, the absolute numbers for any given event remained largely the same (Fig. 4B). This finding probably reflects that any given exon-skipping event is only present in a tiny fraction of the cell population, and compromised RNA surveillance will not substantially increase the population average of a specific splice variant but will increase the number of different events. A total of 55 exon-skipping events were present at one or more copies per cell in at least 1 of 46 samples from different physiological conditions (vegetative growth, stationary phase, quiescence, glucose depletion, heat-shock, and meiosis; Supplemental Table S7). However, even these higher numbers were variable among replicates and were further elevated in nuclear RNA surveillance mutants. We ranked these events based on their reproducibility across the 46 samples and identified only 5 exon-skipping events that exhibited one or more copy in at least 5 samples (Supplemental Table S7). We therefore conclude that exon-skipping events for any given transcript only occur in a small portion of cells in a population. This result further corroborates the view that exon-skipping events reflect splicing errors.

### Unannotated exon-skipping events in human show similar characteristics as in fission yeast

There is ample evidence for alternative splicing in human cells (Wang et al. 2008). Recent findings, however, are raising questions regarding the extent and functional relevance of alternative splicing (Melamud and Moult 2009; Pickrell et al. 2010; Gonzalez-Porta et al. 2013). We therefore tested to what extent exon-skipping transcripts are actively degraded by the exosome also in human cells. To this end, we generated a database containing all theoretical exon-exon junctions for all transcripts, based on human Ensembl annotations (Flicek et al. 2014). Against this database, we aligned human RNA-seq data acquired after siRNA depletions of the following factors: non-catalytic exosome component hRRP40 (Ntini et al. 2013), cytoplasmic hDIS3 homologue (hDIS3L), and cytoplasmic 5’-3’ exonuclease hXRN1, alongside their non-depleted controls (Lubas et al. 2013).

Using a collapsed database (Methods), we identified 30,380 transcripts from 5,366 genes that carried 8,945 unique exon-skipping events. Of these, 2,536 exon-skipping events from 1,903 genes and 10,693 transcripts were present across 8 samples, corresponding to the exon-skipping set annotated in Enseml. However, only ∼26% of these events (671/2,536) were supported by at least 1 skipping-read in each sample. We also identified 16,446 unannotated exon-skipping events from 6,251 genes and 24,009 transcripts (henceforth called novel exon-skipping set; Supplemental Table S8), of which only ∼2.5% (412/16,446) were identified in all samples. These novel exon-skipping transcripts in human were pervasively spliced and supported by a similar number of RNA-seq reads as for the annotated set. Thus, the majority of exon-skipping events in both the annotated and novel sets were not consistently identifiable across the entire dataset.

Intriguingly, the unannotated exon-skipping events showed remarkably similar, although less pronounced, characteristics to those observed in fission yeast (Fig. 5A). Many of both the novel and annotated exon-skipping events appeared stochastic and were only supported by low numbers of diagnostic skipping-reads (∼0.8% of total exon-exon junction reads). Only the novel exon-skipping set, however, showed low but significant accumulation in hRRP40 depleted cells (Fig. 5A; p <2.2e-16, Pearson’s Chi-squared test), while the annotated set exhibited a much weaker and insignificant tendency in the same direction. No enrichment was evident in cells compromised for cytoplasmic RNA surveillance (Fig. 5A).

**Figure 5.**
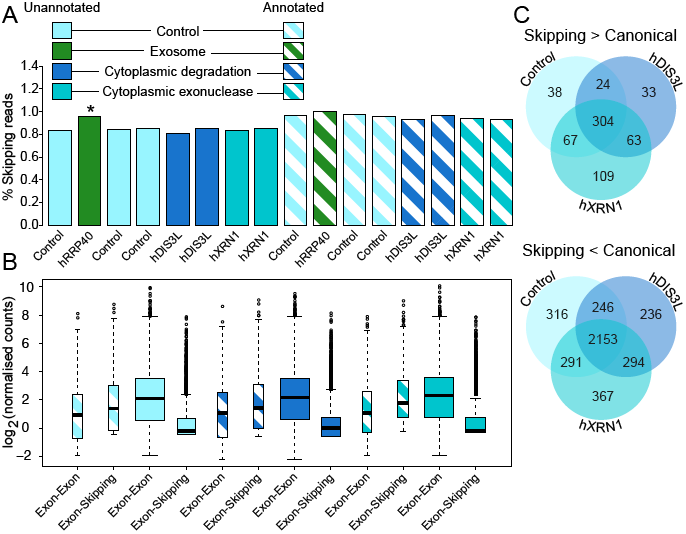
Moderate accumulation of novel exon-skipping in human exosome-depleted cells. (A) Proportion of exon-skipping reads (%) among total exon-exon junction reads in human cells alongside their controls. Exon-skipping events were assigned to annotated and novel sets (striped and solid, respectively; 2,536 and 16,446 exon-skipping events supported by a total of 88,917 and 79,509 exon-skipping reads across all samples, respectively). Significance between different proportions was assessed using the prop.test function in R (p <2.2e-16 for unannotated, Pearson’s Chi-squared test; other comparisons were not significant following Bonferroni correction). (B) Normalised exon-skipping read counts for annotated and unannotated exon-skipping sets (normalised by DESeq package in R; log_2_ scale). Note that the hRRP40 sample was not included due to lack of biological repeats. (C) Up-and down-regulated exon-skipping events relative to corresponding canonical exon-exon junctions in each sample (adjusted p-value <0.05 and fold change of ±2 as determined by DESeq package in R; Supplemental Table S8). Note that annotated and novel exon-skipping sets were combined for simplicity; for corresponding p-values see Supplemental Table S8.

We then compared the abundance of exon-skipping events to the corresponding canonical splicing events for both the annotated and novel exon-skipping sets. On average, novel exon-skipping read numbers were significantly lower than the corresponding canonical exon-exon junction reads in all conditions (Fig. 5B; p <2.2e-16, Wilcoxon Rank Sum test). The annotated exon-skipping events, on the other hand, were significantly more frequent on average, compared to their canonical exon-exon counterparts (Fig. 5B; p <2.2e-16). Overall, across both sets only a small portion of exon-skipping events were consistently higher than the canonical reads in all conditions, whereas the vast majority were significantly and consistently lower (Fig. 5C).

Similar to fission yeast, only 38.6% (6,352/16,446) of the novel skipped exons in human were divisible by 3; thus, the remaining novel exon-skipping events likely interfere with the open reading frame. In contrast, 84.3% of the annotated skipped exons were divisible by 3 (2,138/2,536), suggesting that the majority are likely to maintain the open reading frame. This trend might also reflect a bias in their annotation by automated pipelines.

Taken together, our analysis reveals pervasive but cryptic human exon-skipping transcripts that accumulate to some extent in exosome-depleted cells. These novel exon-skipping events are as ubiquitous as the annotated exon-skipping transcripts. Furthermore these unannotated transcripts are supported by a comparable number of RNA-seq reads, yet they are more stochastic and mostly much lower expressed compared to their canonical counterparts. These findings suggest that some of these novel splice variants may reflect splicing errors that are actively degraded by the exosome. Most annotated exon-skipping transcripts, on the other hand, do maintain their open reading frames and are on average more highly expressed compared to the corresponding canonical transcripts. The findings that the annotated exon-skipping transcripts were often higher expressed and exhibited only marginal accumulation in exosome-depleted cells suggest that they mostly do not reflect splicing errors, and they might show more robust expression in specific developmental stages, tissues or conditions not sampled here.

### Splicing efficiency in RNA surveillance mutants

Accumulation of pre-mRNA is a hallmark of cells with compromised RNA surveillance (Bousquet-Antonelli et al. 2000; Lemieux et al. 2011). To evaluate the extent of pre-mRNA accumulation, we computed the splicing efficiency (SE) as the ratio of mRNA to pre-mRNA for each intron (Fig. 1A). Splicing efficiency in fission yeast was significantly lower in exosome mutants compared to wild-type, and it also gradually decreased as meiotic differentiation progressed (Supplemental Fig. S6A). Regulated splicing, where splicing efficiency is dramatically increased during meiosis is a well-documented phenomenon in fission yeast (Averbeck et al. 2005; Wilhelm et al. 2008; Chen et al. 2011). Accordingly, we also detected the introns whose splicing efficiency increased during meiosis (Supplemental Fig. S6B). As previously documented (Livesay et al. 2013; Bitton et al. 2014), splicing efficiency was also significantly lower in splicing mutants. Splicing efficiency was highly correlated with gene expression, indicating that introns in highly expressed genes are more efficiently spliced than those in lowly expressed genes (Supplemental Fig. S7). This intriguing finding confirms previous data (Wilhelm et al. 2008) and may reflect the tight coordination between transcription and splicing machineries. In contrast to fission yeast, splicing efficiency was not globally affected in human exosome-depleted cells (Supplemental Fig. S8). This finding is in accordance with a recent study reporting retention of only a few major-spliceosome (U2-type) introns in nuclear exosome knock-down cells (Niemela et al. 2014).

### Pervasive distribution of cryptic splice sites in fission yeast

Exon-skipping represents only one mechanism by which alternative splice variants could be generated; additional mechanisms are intron retention and alternative 5’-and 3’-splice donor and acceptor sites. Recent RNA-seq studies reported numerous novel introns in budding yeast (Volanakis et al. 2013; Kawashima et al. 2014) and fission yeast (Awan et al. 2013; DeGennaro et al. 2013; Lee et al. 2013). We developed an exhaustive search algorithm to partition the genome based on all possible splice donor and acceptor di-nucleotide combinations, rather than considering only the consensus 5’-(GU) and 3’-(AG) splice sites (Supplemental Fig. S9). By doing so, we generated a database containing all junctions that bridge all possible annotated or novel introns. Against this database, we then aligned all RNA-seq reads that could not be mapped to the genome or annotated transcriptome. A clear enrichment for consensus 5’-and 3’-splice signals was evident (97.7% of novel events), suggesting that splicing of these novel events is mediated by the major spliceosome (Supplemental Fig. S10). Although a few non-consensus splice site signals were uncovered, they were of lower frequency than the expected ∼5% false identification. Considering only consensus 5’-and 3’- splice signals, we identified a total of 5,720 unannotated splicing events (henceforth called novel introns) that were supported by 1,165,427 unique junction reads. We also identified a set of 307 known introns that were supported by 657,527 unique junction reads (Supplemental Table S9). These known introns were picked up because some mRNA transcripts were mis-annotated or omitted from the Ensembl transcriptome file used here (release 13); their identification therefore provided a positive control and corroborated our approach.

The 5,720 novel introns were distributed predominantly within protein-coding genes (∼61%), but also in non-coding RNAs and intergenic regions (∼38% and ∼1% overlapping accessions, respectively; Supplemental Fig. S11). Interestingly, 967 protein-coding genes with novel introns have not been known to be spliced. Novel introns displayed a similar length distribution to the annotated set of 5,361 introns, with a marginal tendency to be longer (Supplemental Fig. S12). They also had near identical splice-signal consensus sequences (Supplemental Fig. S13), located in comparable distances relative to the 3’-ends of introns. However, novel introns exhibited higher GC content than annotated introns (Supplemental Fig. S14).

Taken together, these findings suggest that the novel introns represent both *bona-fide* introns as well as cryptic introns. This interpretation was corroborated by the skewed distribution of the number of samples in which novel introns appeared (Fig. 6A), and by the distribution of the number of supporting junction reads (Fig. 6B). Most novel introns only appeared in a small number of samples and were supported by low numbers of junction reads, and these introns are likely to reflect splicing errors. Much fewer introns were identified in most of the samples and were supported by a high number of reads, and these introns are likely real, novel introns. There was a high correlation between the number of unannotated splicing events identified and sequencing depth, implying that most of these introns are simply rare, aberrant events (Fig. 6C). As for exon-skipping, these unannotated splicing events tended to accumulate in cells with defective nuclear degradation, and also in cells with compromised transcription regulation due to the lack of functional Spt6 (Fig. 6C, indicated by arrows). We tested 30 of the novel introns by RT-PCR: 29/30 displayed the expected size, and 25/30 were also validated by Sanger sequencing (Supplemental Fig. S15; Supplemental Table S10).

**Figure 6.**
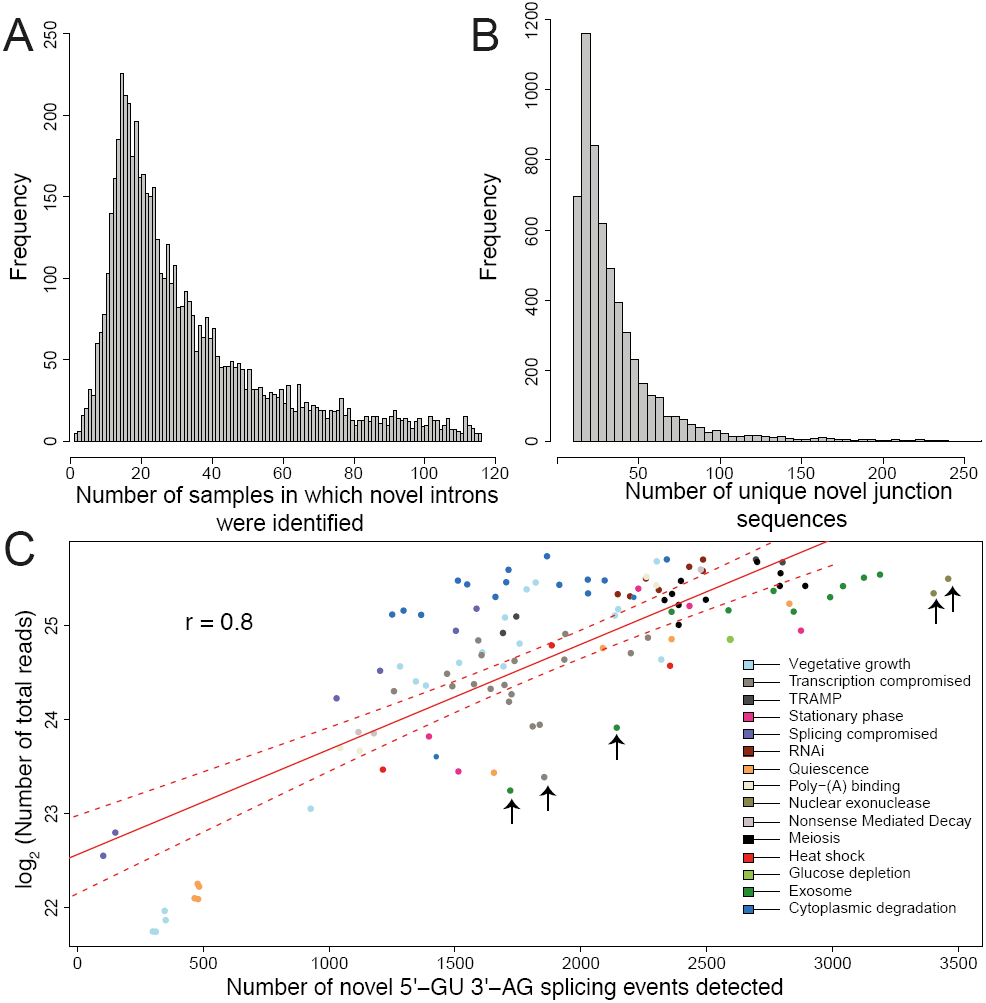
Distribution of novel introns with respect to (*A*) samples in which they were identified, and (*B*) the sum of their supporting unique sequences across the entire junction and the entire dataset. A total of 5,720 unannotated splicing events are shown. (*C*) Correlation between the samples’ sequencing depth and number of novel 5’(GU)-3’(AG) splicing events; r: Pearson’s correlation coefficient; red line: fitted regression; dotted red lines represent 0.95 confidence levels. Each dot represents a sample (116 in total), grouped according to cellular functions and conditions. Note that a given splicing event could be identified in multiple samples. Arrows highlight accumulation of cryptic events when degradation or transcription regulation is compromised (e.g. *nmt1-dis3* and *dhp1-ts* or *spt6-ts).*

Expression profiles of selected introns highlighted the differences between novel introns that were missed by current annotation and those that were more cryptic (Supplemental Fig. S16).

We conclude that the novel introns represent both a minority of novel, *bona-fide* introns that have been missed by prediction algorithms and a majority of inefficiently spliced, cryptic introns which show similar characteristics as the exon-skipping events analyzed above. These findings reveal that splicing of the fission yeast transcriptome is much more pervasive than appreciated, but most of these novel events might reflect splicing errors without cellular function.

## Discussion

Splicing is pivotal for gene expression, and mistakes during this process can be detrimental. Alternative splicing has been considered mainly as a mechanism by which cells augment their molecular toolbox at the proteome level, with >90% of human genes potentially producing multiple transcripts from a single locus (Wang et al. 2008; Wang and Burge 2008). Here we present data highlighting that many alternative splice variants are not necessarily functional. Using a systematic analysis of RNA-seq data from fission yeast, we conclude that widespread but low frequency exon-skipping events reflect splicing errors that are actively degraded within the nucleus. Our analysis of human data indicates that widespread, cryptic exon-skipping events also occur in multicellular eukaryotes, although many annotated global ESRs show approximately 30-fold higher frequencies than in fission yeast and probably represent *bona-fide* transcript isoforms. These results raise the possibility that alternative splicing has originally emerged as a by-product of cellular splicing errors, which have then been co-opted during evolution for specialized protein functions in complex eukaryotes. This hypothesis is supported by two recent studies showing that alternative splicing signature is species-specific, and significantly differs even between closely related species (Barbosa-Morais et al. 2012; Merkin et al. 2012).

We have previously shown that exon-skipping is not a consequence of bioinformatics noise, by validating a subset of exon-skipping events using gene-specific RT-PCR and identifying 82 exon-skipping lariat-reads in de-branching deficient *dbr1*Δ cells (Bitton et al. 2014). The following pieces of evidence, however, indicate that exon-skipping transcripts in fission yeast represent splicing errors which might have no functional importance. First, exon-skipping is a rare and erratic phenomenon, supported by only few RNA-seq reads relative to reads supporting regular splicing. Second, exon-skipping events are randomly scattered throughout the transcriptome, showing no obvious dependency on intron or exon length, on expression levels, or on environmental conditions (except a slight increase during late meiotic stages); however, they are positively correlated with the number of splicing reactions a given gene undergoes, in line with previous reports (Pickrell et al. 2010). Third, exon-skipping transcripts are not only much less abundant compared to their canonical counterparts, but also accumulate in cells with impaired nuclear RNA surveillance and during late meiotic differentiation when RNA surveillance, but not the splicing machinery, might be down-regulated. Fourth, they show low cellular quantities below 1 RNA copy/cell on average for a given transcript in a population; such low levels are diagnostic for actively repressed transcripts that have no function under the given condition (Marguerat et al. 2012). Fifth, exon-skipping events show no enrichment above random expectation to preserve the open reading frames, and the majority of these events are therefore expected to lead to premature stop codons. Nevertheless, we cannot exclude that some of the exon-skipping events identified become functionally relevant during specialized conditions not investigated here, e.g. in specific stages of the cell cycle.

The low copy numbers for exon-skipping transcripts in fission yeast indicate that there are only very few cells in a population that produce a given splice variant. In the absence of nuclear RNA surveillance, we observed a dramatic increase in the diversity of exon-skipping events, but no corresponding increase in the copy numbers of specific splice variants. These findings corroborate the interpretation that exon-skipping represents stochastic splicing errors that vary from cell to cell and that are kept in check by RNA quality control. These findings can also explain why the number of exon-skipping events do not correlate with the expression level of specific transcripts, but do correlate with overall sequencing depth. Among the pervasive set of exon-skipping events in fission yeast, however, 111 exons were more prone to be skipped, although they also appear to represent splicing errors. We could not pinpoint any common characteristic or DNA sequence features among these exons that might explain why they are much more frequently skipped than other exons.

A focused analysis of four loci in budding yeast uncovered the involvement of Rat1/Dhp1 and the nuclear exosome in degrading transcripts that carry exon-skipping information (Egecioglu et al. 2012). Here we show that exon-skipping events are degraded by the same two surveillance pathways in the distantly related fission yeast, and, to a lesser extent, the RNA exosome also degrades exon-skipping transcripts in human cells. In fission yeast, we also observed a relatively low, but significant, accumulation of exon-skipping transcripts in cells with mutations in other RNA surveillance pathways, like TRAMP and NMD. These data suggest that several different RNA surveillance mechanisms can recognize erroneous exon-skipping events and back each other up (Sayani and Chanfreau 2012), although the nuclear exosome and Rat1/Dhp1 play dominant roles. It is not clear how these two pathways can recognize aberrantly spliced transcripts. Surprisingly, NMD seems to play only a minor role in degradation of aberrantly spliced transcripts. Recent reports in budding yeast show that inefficient splicing could modulate gene expression via NMD (Kawashima et al. 2014) or SMD (Volanakis et al. 2013). Similarly, widespread intron retention in mammalian transcriptomes has recently been shown to tune gene expression (Braunschweig et al. 2014). However, in our data relative changes in abundance of specific splice variants between mutant and wild-type samples were not accompanied by changes in corresponding transcript levels (data not shown). We therefore conclude that exon-skipping is not widely used to modulate gene expression in fission yeast.

Our exon-skipping data, together with the absolute copy numbers of transcripts per cell, enabled a more accurate estimation of splicing errors, revealing that many errors are masked by the activity of RNA surveillance mechanisms. In fact, the population’s error rate could be even higher in the absence of both pathways or of alternate RNA surveillance pathways that could take over in the absence of the nuclear exosome or Rat1/Dhp1.

Exon-skipping transcripts increased during late stages of meiotic differentiation. The exosome is known to ensure timely expression of meiotic transcripts via the Mmi1 system (Harigaya et al. 2006), and many meiotic transcripts show regulated splicing (Averbeck et al. 2005). However, our data do not support any functional role of exon-skipping during meiosis. Even the increased levels of exon-skipping transcripts are still at very low abundance relative to the corresponding conventionally spliced transcripts. We also observed a gradual decrease in splicing efficiency and a decrease of nuclear-exosome and *dhp1* transcripts during sexual differentiation. These results may suggest that the increased exon-skipping transcripts in late meiosis result from lowered nuclear surveillance activity. The purpose, if any, for this down-regulation of RNA surveillance during meiosis remains to be elucidated. It has been reported that Rrp6 is degraded during meiosis in budding yeast which plays a regulatory role for meiotic progression (Lardenois et al. 2011). However, a recent study has shown that the exosome remains active during budding yeast meiosis (Frenk et al. 2014).

Although a high degree of correlation was observed between gene expression and splicing efficiency, there was no strong effect for any of the CTD mutants of RNA polymerase II on splicing efficiency or exon-skipping. However, the global decline in splicing efficiency observed in exosome mutants and during meiosis fits well with the model of substrate-competition between the exosome and spliceosome (Bousquet-Antonelli et al. 2000; Lemieux et al. 2011).

RNA splicing and surveillance pathways are largely conserved between fission yeast and human (Käufer and Potashkin 2000). Interestingly, analysis of exon-skipping events in human data exposed several trends that resemble those in fission yeast. First, we uncovered numerous cryptic, lowly expressed exon-skipping transcripts. Second, these novel transcript isoforms were only supported by a low number of reads, as were those associated with annotated exon-skipping events; only a small fraction of exon-skipping events in either annotated or novel sets were consistently detectable across the entire dataset. Third, the novel exon-skipping transcripts displayed a slight accumulation in hRRP40 depleted cells, and the majority of these transcripts were significantly lower expressed compared to their canonical counterparts. These observations are in accordance with a recent report revealing that a single dominant transcript is present for most human genes across a wide range of tissues and cell lines (Gonzalez-Porta et al. 2013).

Several technical or biological factors could explain the moderate accumulation of exon-skipping events and pre-mRNA transcripts in human cells depleted for the nuclear exosome compared to fission yeast: 1) siRNA might not have completely abolished protein function compared to yeast gene knock-outs, although efficient depletion has been reported (Lubas et al. 2013; Ntini et al. 2013); 2) redundancy of RNA surveillance pathways; and 3) relatively shallow sequencing depth. While the first two factors might directly affect accumulation of aberrantly spliced and pre-mRNA transcripts, the latter would also account for the poor and inconsistent identification of lowly expressed transcripts. In any case, our data do not support the view that all human exon-skipping events represent *bona-fide* transcripts that are functionally relevant. Their low and erratic expression, which in the case of unannotated transcripts is dwarfed by conventionally spliced transcripts of the same gene, along with their likelihood to alter the open reading frame, strongly suggest that much of these exon-skipping events represent splicing errors under the condition analyzed here, and likely also under other tested conditions (Gonzalez-Porta et al. 2013). We show that exon-skipping transcripts in fission yeast are largely expressed at below 1 copy per cell. Single-cell analyses of mouse transcriptomes revealed extensive heterogeneity in splicing patterns between individual cells, with a strong bias towards a single splice variant in any given cell (Shalek et al. 2013). These findings further support that alternative splice variants are often produced erroneously.

Our exhaustive search for cryptic splice sites has doubled the number of splicing events in the well-studied fission yeast. While the robust set of novel introns demands genome-wide reassessment of current annotation, the larger set of lowly expressed, inefficiently spliced and unstable introns further highlights that splicing errors are far more common than previously recognized. Similar issues have been documented also in human cells (Melamud and Moult 2009; Pickrell et al. 2010). A recent interrogation of the translational landscape in fission yeast has revealed only a single case of intron retention, yet numerous examples of incorrect annotation (Duncan and Mata 2014), supporting our view that the vast majority of novel splicing events represent splicing errors. Similar to exon-skipping events, we show that these inefficiently spliced products also tend to accumulate in cells with compromised nuclear RNA surveillance pathways. As it is impossible to systematically distinguish unannotated novel introns from aberrantly spliced introns, we cannot exclude the possibility that some introns are in fact alternatively spliced under certain conditions.

Our analyses, together with diverse observations made by several other studies (Melamud and Moult 2009; Pickrell et al. 2010; Gonzalez-Porta et al. 2013; Shalek et al. 2013), refine the dogma that alternative splicing in eukaryotes is limited to the production of functionally relevant transcripts. Even in human cells, a considerable fraction of alternative splice variants could simply reflect splicing errors, which in turn is actively targeted by RNA surveillance mechanisms.

## Methods

### Strain list, RNA isolation and sequencing

The genetic background, growth conditions, type of RNA-seq (i.e. poly-(A) enriched, total RNA or ribosomal depletion), and the sources of all *S. pombe* strains used in this study are specified in Supplemental Table S1. For poly-A enriched libraries, RNA extraction, library preparation and sequencing protocols were as described be Lemieux et al. (2011), for total RNA as described by Marguerat et al. (2012), for ribosomal-depleted libraries as described by Bitton et al.(2014). Publically available datasets were downloaded from GEO, SRA or ArrayExpress repositories (accessions indicated in Supplemental Table S1). For some paired-end datasets (Livesay et al. 2013; Schwer et al. 2014) only the 5’-end of the read was used throughout, while for another paired-end dataset (Rhind et al. 2011), both ends of the read were used exclusively for cryptic intron discovery. Human datasets were retrieved from GEO: GSE48286 (Ntini et al. 2013; Lubas et al. 2013). Only the 5’-ends of paired-end reads were used in the human transcriptome analysis.

### Genome level alignments and annotation

Sequence reads of ‘X’ base length (see Supplemental Table S1 for exact read length of each RNA-seq experiment) originating from each sample were aligned using Bowtie 0.12.7 (Langmead et al. 2009) to the *S. pombe* genome sequence (Ensembl *S. pombe*, Build EF1, release 13. Flicek et al. 2014) and to the corresponding exon-exon junctions database. Up to 3 base-pair mismatches were allowed. Reads that matched multiple loci were removed from further analysis, and the resultant alignment files were processed to generate ‘pile-ups’ against each chromosome (for total number of mappable reads in each sample, see Supplemental Table S1). Importantly, throughout our exon-skipping analysis we used only canonical and non-canonical exon-exon junction reads (Fig. 1A) and reads that straddle the exon-intron boundaries (Fig. 1A), while the remaining reads were ignored. Furthermore, reads that could not be mapped to the genome or transcriptome were used for cryptic intron analysis. However, for correlations of skipping frequency or splicing efficiency with gene expression (Supplemental Figs. S3, S4 and S7), normalised expression levels were calculated as described (Bitton et al. 2014). Human datasets were analysed using the same criteria but searched against Ensembl human annotation (release 70; Flicek et al. 2014), combined with a reduced exon-exon junction database (see sub-section below).

### Exon-exon junctions and exon-skipping databases

Searches were performed against the genome sequence combined with a database of all possible exon-exon junction sequences that could be generated from Ensembl *S. pombe* annotation (release-13). To ensure that an ‘X’-base read is mapped to a splice junction (see Supplemental Table S1 for exact read length), only the last (’X’-6) bases of the first exon and the first (’X’-6) bases of the second exon were considered (if the exon exceeded length ‘X’-6). In this way, reads that overlapped a junction by less than 6 nucleotides were excluded. Reads that matched to more than one junction or elsewhere in the genome were also discarded.

Human datasets were analysed with the same criteria, but using Ensembl human annotation (release 70; Flicek et al. 2014). Since a typical human gene has multiple transcripts and multiple overlapping exons, for each transcript all possible combinations of exons were initially considered (5’-3’) to create all possible exon-exon junctions. Thereafter, the database was collapsed based on unique junction sequences to avoid redundancy (all junction information was recorded, see example in Supplemental Table S8 “Complete_junction_details” column). Furthermore, to avoid splitting the database due to its enormous size, we first aligned the RNA-seq data against the exon-exon junction database alone and pulled out all alignments that matched our criteria (retaining hits with up to 3 mismatches and a unique alignment, as before). We then constructed a reduced exon-exon junction database of all positive hits in all samples and combined it as an additional chromosome to allow simultaneous search together with the genome sequence, while the same mapping criteria applied.

### Cryptic splice site analysis

To identify cryptic splice sites independently of prior splice-site knowledge, we first divided the fission yeast genome into six batches (5’-3’ direction), based on all annotated genes, as follows. In each batch, we stored a fraction of the genome containing different gene sequences ±300 bp upstream and downstream to ensure identification of introns within UnTranslated Regions (UTRs), and any intervening sequences (i.e., intergenic regions). Thereafter, we partitioned each region within each batch based on all possible splice donor and acceptor di-nucleotide combinations (i.e. 4^2^ × 4^2^ = 256 combinations; Supplemental Fig. S9). By doing so, we generated a database containing all possible introns (a minimum size of 10 bp in length) around which splice junctions were constructed, as follows. Our diverse dataset included RNA-seq reads of varying lengths (49-76 bp), to maximise read mapping we created junctions around putative introns using 57 bp upstream and 57 bp downstream of their flanking sequences (or entire flanking sequence if shorter than 57 bp). Thereafter, for each batch we stored, as a single database, all junctions produced from a given splice donor and acceptor combination. Then, using Bowtie, we aligned to each database all RNA-seq reads that could not be mapped to the genome or the annotated transcriptome (33 samples), but this time retained only unique matches without tolerating any mismatches. The remaining 83 samples were aligned only to databases produced from the consensus 5’-(GU) and 3’-(AG) splice donor and acceptor sites. Thereafter, read alignments from all databases, di-nucleotide combinations and samples were combined into a single file. Using customised Perl and R scripts, we filtered out non-unique matches, or those from rRNA regions in both orientations as well as reads with an overhang of less than 6 bp across the junction. To further minimise false positive mappings, we calculated the False Discovery Rate (FDR) based on results obtained from alignments of 33 samples that were searched against all possible di-nucleotide combinations. To do so, we considered all combinations other than 5’-GU and 3’-AG splice donor and acceptor sites as random matches. Then, we calculated the FDR as follows: FDR= (number of random junctions)/(number of 5’-GU and 3’-AG junctions + random junctions), while the number of unique sequence reads starting at different locations along the junction was used as the varying cutoff. FDR analysis revealed that only ≤5% random matches were retained in the reported list if a given junction was supported by >13 unique sequences across 33 samples (Supplemental Fig. S10). This threshold was then applied to junctions obtained from all 116 transcriptomes. Under this threshold, we identified 6,238 putative junctions (5’-GU - 3’-AG junctions only). However, due to the ±300 bp upstream and downstream gene flanking sequences that were used for our database, there was a low degree of redundancy. Therefore, 5,899 novel splicing events led to the identification of 5,720 novel introns (supported by 1,165,427 unique junction reads), while the remaining 339 events corresponded to 307 known introns (supported by 657,527 unique junction reads).

### Comparison of novel and annotated introns

The GC content of both the novel and annotated introns sets was determined using the ‘geecee’ function within the European Molecular Biology Open Software Suite (EMBOSS) (Rice et al. 2000). Branch-site predictions were performed using FELINES (Drabenstot et al. 2003). Two FASTA files containing 5,686 novel and 5,283 annotated introns with canonical splice donor and acceptor sites (GU and AG, respectively) and of length greater than 20 bp were analysed using FELINES with default settings. Consensus branch-site, 5’ and 3’ sequence logos were plotted using WebLogo (Crooks et al. 2004).

## Data Access

RNA-seq datasets produced in this study are available from the European Nucleotide Archive (ENA) under accession number PRJEB7379.

## Acknowledgments

We thank Tristan Clark and David Gregory for their help with high-performance computing, and Beate Schwer for comments on the manuscript. This work was supported by a Wellcome Trust Senior Investigator Award (grant # 095598/Z/11/Z) to J.B. and the Danish National Research Foundation (grant DNRF58) to T.H.J.

### Author Contributions

DB designed the study and conducted the bioinformatics analysis. DJ, SM assisted with algorithm design. DB and DJ designed the statistical approach. CR and DE experimentally validated novel introns. GS characterised novel introns using FELINES and did the artwork. CC and JM produced the meiotic time-course data. ML and THJ assisted with designing the analysis of the human dataset. DB, SA, MM, SC, YC, SM, FB prepared the different RNA-seq libraries. DB and JB conceived the study and wrote the manuscript.

### Disclosure Declaration

The authors declare that there are no conflicts of interest.

